# Class-specific sensing of HIV-1 antigens by the B cell antigen receptor depends on the CH1 domain

**DOI:** 10.1101/2023.12.13.571545

**Authors:** Yaneth Ortiz, Kara Anasti, Advaiti K. Pane, Ken Cronin, S. Munir Alam, Michael Reth

**Author notes:** **Author contributions:** Y.O. and M.R. designed the research. Y.O. performed the experiments. K.A., A.K., K.C., and S.M.A. characterized the HIV-1 Env proteins and selected the antigens for the study. M.R. Y.O. and S.M.A. wrote the paper.

## Abstract

How different classes of the B cell antigen receptor (BCR) sense viral antigens used in vaccination protocols is poorly understood. Here we study antigen binding and sensing of Ramos B cells expressing BCRs of either the IgM or IgG1 class with a specificity for the CD4-binding-site of the envelope (Env) protein of the human immune deficiency virus-1 (HIV-1). We find that, in spite of their identical antigen binding site, the two BCR classes differ drastically from each other in that the IgM-BCR and IgG1-BCR bind preferentially to monovalent and polyvalent antigens, respectively. By generating an IgM/IgG1 chimeric BCR we found that the class-specific antigen-sensing behavior can be transferred with the CH1γ domain from the IgG1-BCR to the IgM-BCR. Our results indicate that the class-switching process not only results in the production of antibody classes with different effector functions but also alters the antigen sensing of secondary B lymphocytes. These findings suggest that antigen valency in existing vaccination protocols should be modified and altered between primary versus secondary (booster) immunization.

## INTRODUCTION

The acquired immunodeficiency syndrome (AIDS) caused by the human immunodeficiency virus (HIV-1) is a major health crisis worldwide (1). Although drugs that suppress virus replication in infected persons have been developed, the spread of HIV-1 would be most efficiently prevented by the development of an anti-HIV-1 vaccine (2). Unfortunately, due to the high mutation rate of this retrovirus, this is a challenging task. One strategy in HIV-1 vaccine development is to employ viral epitopes that cannot be easily altered by the virus without losing its ability to infect human T cells. The CD4 binding site (CD4bs) within the trimeric envelope (Env) protein of HIV-1 is such an epitope.

Many monoclonal antibodies (mAbs) that target this conserved epitope have been generated and it was shown that they have broad neutralization capacity against the majority of HIV-1 strains isolated from infected humans(3–5). These broadly neutralizing antibodies (bNAbs) have also been found in AIDS patients. It has been shown that passive transfer of bNAbs to non-human primates can provide sterilizing protection against chimeric simian/human immunodeficiency viruses (6). Thus, it is widely expected that vaccines inducing sustained titers of bNAbs would protect humans against HIV-1 (6, 7).

Despite many trials conducted over the last decades, the development of an efficient vaccine that induces the production of bNAbs in humans has not been achieved. During its expansion, HIV-1 counteracts an immune blockade by generating Env protein with immunodominant epitopes outside the CD4bs or with mutations that partially block access to the CD4bs (8, 9). The current anti-HIV-1 vaccine development is thus focused on fragments or altered forms of the HIV-1 Env protein carrying an accessible and immunodominant CD4bs. The study of the humoral immune response against the CD4bs epitope is supported by the generation of a large panel of anti-CD4bs bNAbs used in binding studies of altered Env proteins. CH31 is a V_H_1-2 bnAb of the VRC01 CD4bs family (CD4-mimic) for which the affinity and kinetics of binding to diverse HIV-1 Env-derived proteins have been determined (10– 13). The VRC01 family presents a key model as these bnAbs are the most broadly neutralizing and their precursors are populated at a relatively higher frequency (14).

What is less well known is how the variant Env antigens are sensed by different classes of the B cell antigen receptor (BCR). The IgM-class BCR (IgM-BCR) is first expressed on immature B cells while mature naïve B cells co-express two BCR classes, namely an IgD-BCR and IgM-BCR. After their first activation, B cells can undergo a class switch and express either an IgG-, IgA-, or IgE-BCR (15). In their basic structure all classes of the BCR are similar and form a complex between the membrane-bound immunoglobulin (mIg) molecule and the CD79a/CD79b (Iga/Igb) heterodimer mediating antigen-binding and signal transduction, respectively (16–18). The mIg molecule is assembled in the endoplasmic reticulum (ER) and forms a homodimer of two identical heavy chains (HC) and two identical light chains (LC). Most mammals have two types of light chains namely kappa (κ) and lambda (λ) with a different gene organization and distinct sequences of their Cκ and Cλ domains. The antigen binding site (also called paratope) is formed by the variable domains (VH/VL) of each HC: LC pair. Thus, a mIg molecule carries two identical antigen binding sites. The recently published cryo-EM structures of the IgM-BCR and IgG-BCR show the molecular details of the asymmetric mIg: CD79a/CD79b complex assembly but they do not reveal the activation mechanism of these receptors (19–21). A recent analysis of these structures suggests that BCR activation is accompanied by dissociation and reassociation processes (22). How antigen is sensed by naïve normal B is difficult to study as their antigen specificity is highly variable and, in most cases, unknown. However, with the advent of CRISPR-Cas technology, it is now feasible to generate antigen-specific human B cell lines. In particular, the human Burkitt lymphoma B cell line Ramos is a valuable model for antigen sensing studies (23). Recently, Ramos B cells were generated expressing a IgM-BCR with the VH/VL domains of the CH31 bNAb CH31 (11). It was found that the CH31 IgM-BCR on these cells preferentially sense Env proteins that display a high on-rate of binding to bNAb CH31. We here compared the binding of a CH31 IgM-BCR and CH31 IgG1-BCR to different forms of the Env antigen and found that in spite of their identical CH31 paratope, these antigen receptors differ drastically in their binding characteristics. Specifically, the IgG1-BCR binds preferentially polyvalent antigens and this feature can be transferred with the CH1γ domain onto the IgM-BCR.

## RESULTS

### Generation of HIV-1 (CH31)-specific IgM-BCR or IgG1-BCR Ramos B cells

For the present study, we used the BCR-negative, MDL-AID KO Ramos B cell line(23) and pMIG retroviral transfection vectors to generate CH31 IgM-BCR or CH31 IgG1-BCR expressing Ramos B cells (Figure 1A). After sorting the BCR-positive cells, the level of BCR expression was evaluated by flow cytometry using anti-human κLC antibodies (Figure 1B). Surprisingly, we see that the retroviral transfected Ramos B cells display a 5-fold higher IgG1-BCR than IgM-BCR expression. This class-specific expression pattern was seen in two independently generated mIgG1 or mIgM Ramos B cell transfectants. The staining of the IgM-BCR- or IgG1-BCR expressing Ramos B cells with another anti-BCR antibody, namely that directed against the human CD79b signaling subunit confirmed that the IgG1-BCR is significantly more expressed on the Ramos B cell surface than the IgM-BCR. This result suggests the existence of a class-specific regulation for the export and/or stable expression of the BCR on the B cell surface.

**Figure 1.**
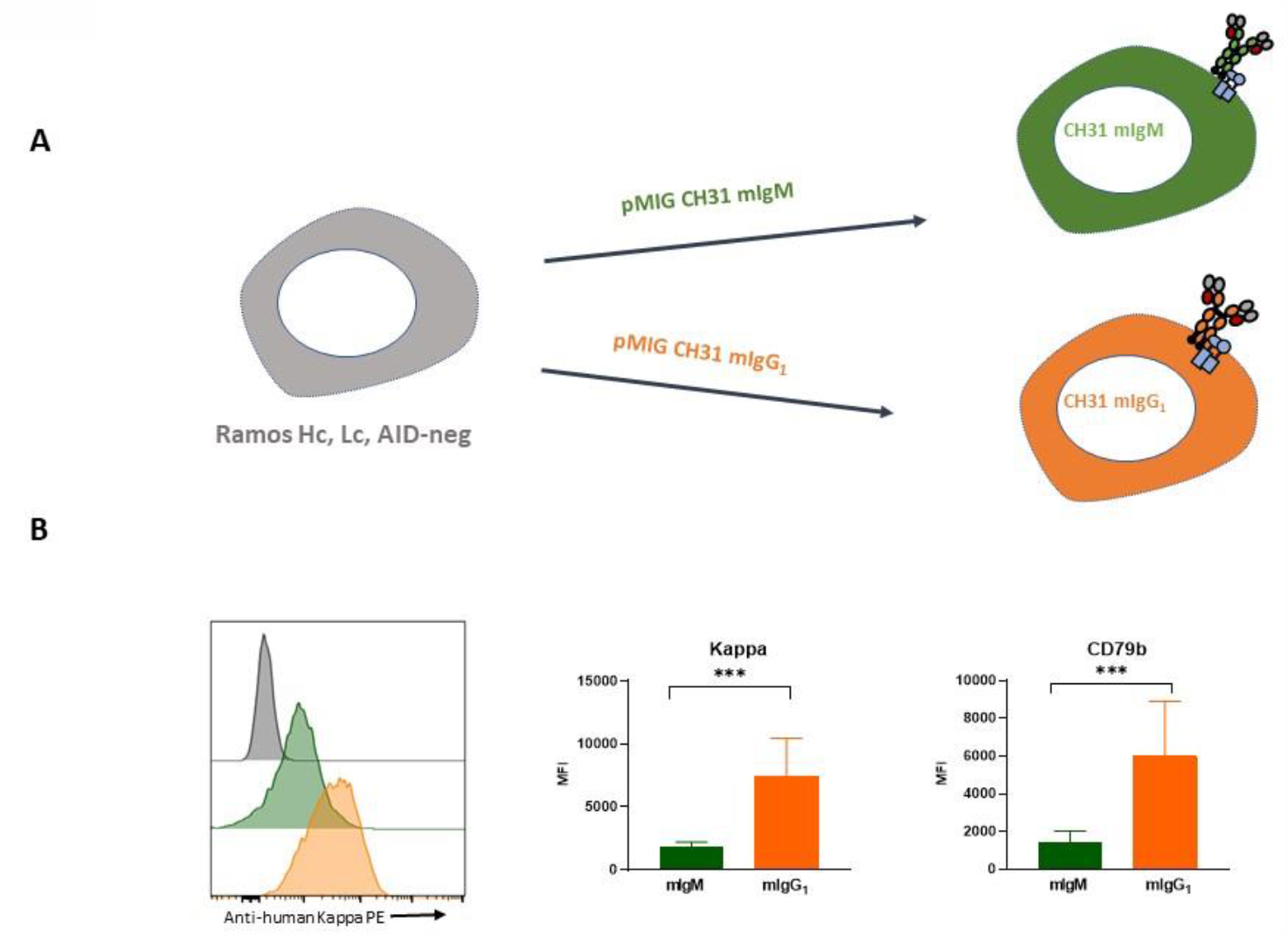
Generation of Ramos B cells with CH31-specific IgM and IgG1 isotypes. **(A)** Strategy for the generation of Ramos B cells with the CH31 bNAbs specificity. Cells were generated on Ramos heavy chain, light chain, AID Knock Out cell line (MDL AID), by retroviral transduction with pMIG plasmids carrying both the heavy chain (IgM or IgG_1_) and the Kappa light chain separated by a P2A sequence. **(B)** Expression of BCR components on CH31 IgM-BCR and IgG1-BCR cell lines, flow cytometry staining for the human κ light chain and CD79b (Igβ) on the IgM (green) and IgG1 (orange) cell lines, MDL AID KO cells (gray histogram) shown as control. PE, phycoerythrin. MFI, median fluorescent intensity, Mann-Whitney test, *** P<0.001

### Phenotype of the IgM-BCR- or IgG1-BCR Ramos B cells

We previously have shown that different classes of the BCR are localized together with other surface proteins inside class-specific protein nanodomains and at distinct topographic locations (24, 25). To test whether or not the IgM-BCR or IgG1-BCR Ramos B cells differ in the expression of other B cell surface proteins we stained these cells with antibodies directed against 29 different B cell surface markers. To directly compare the expression of each of these marker proteins on the surface of the two Ramos cell lines we employed a color-based barcoded flow cytometric assay that we have previously described(26). This analysis confirms that BCR components such as the κLC or the CD79b signaling subunit are expressed in higher amounts on IgG1-BCR than on the IgM-BCR Ramos B cells (Figure 2). Apart from this, the IgG1-BCR Ramos B cells also display a higher expression of the BCR coreceptor CD19 and signal regulator CD45R as well as the tetraspanin CD53 and the chemokine receptor CXCR4 (Figure 2B). The higher expression of this B cell surface marker on the IgG1-BCR Ramos B cells could be either connected to the higher BCR expression and/or to the expression of a different mIg class of these cells (see discussion).

**Figure 2.**
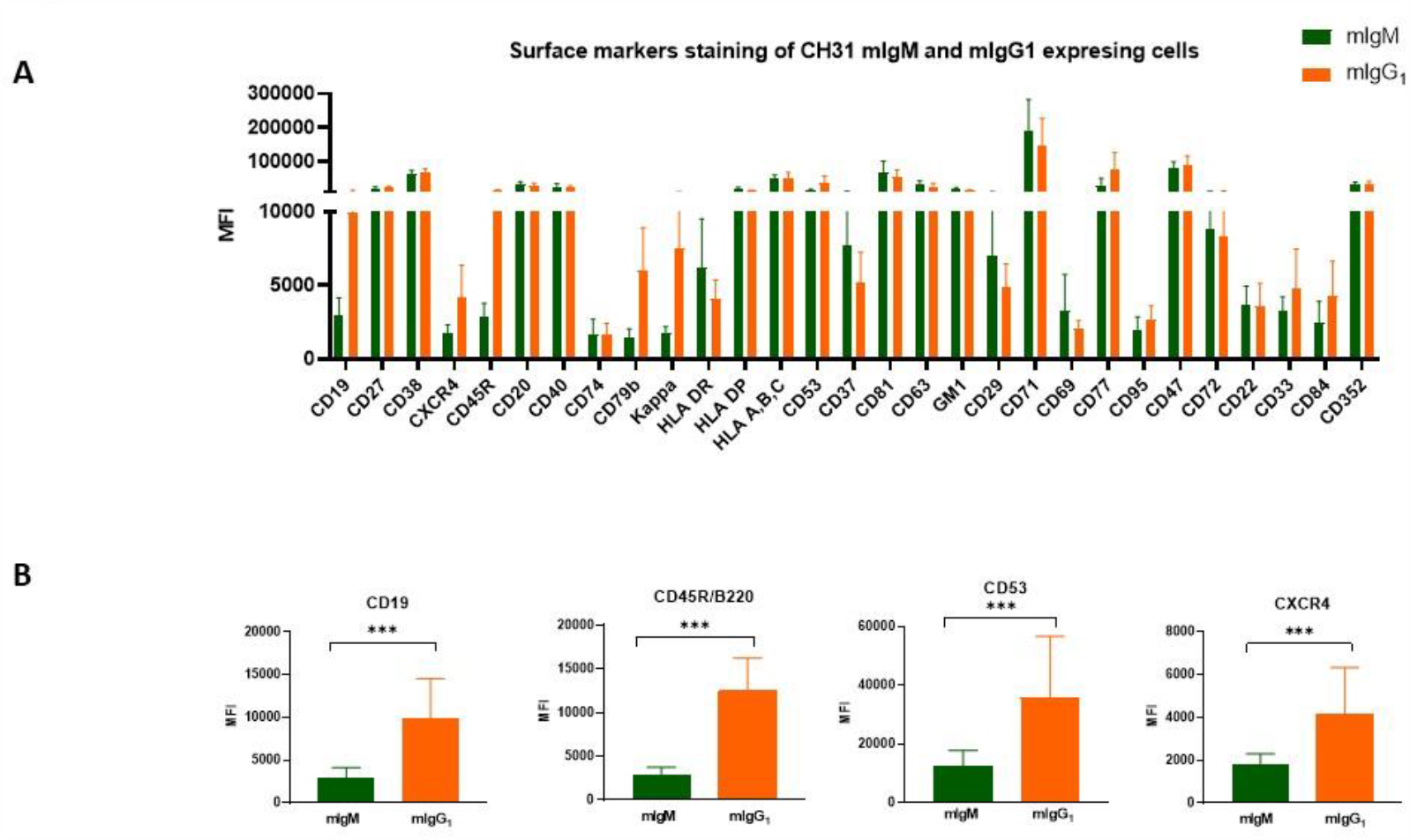
Barcode phenotype of CH31 IgM-BCR and IgG1-BCR cells. **(A)** Comparison of the phenotype of IgM-BCR and IgG1-BCR CH31 cells, cells were barcoded and stained with a panel of antibodies directed against an array of surface proteins (see materials and methods and the antibody table list), shown is the median fluorescence intensity (MFI) and standard deviation (SD) of 29 markers that highlights the differences among IgM-BCR (n=16) and IgG1-BCR (n=12) cell lines. **(B)** B cell surface markers significantly differ among IgM-BCR and IgG1-BCR cells (excluding BCR components). Mann-Whitney test, *** P<0.001.

### Antigen binding characteristics of the CH31 IgM-BCR- or IgG1-BCR-expressing Ramos B cells

Due to the enormous diversity of their V gene repertoire, most ex vivo isolated B cells have an unknown antigen specificity and thus the binding behavior of B cells to their cognate antigen and its variants is not well studied. The MDL-AID KO Ramos B cell system allows one to generate and study human B cells with a defined antigen specificity (27, 28). One of the best-studied B cell antigens is the HIV-1 spike Env protein. Many bNAbs have been described against the HIV-1 spike. The affinity and binding kinetics of these anti-HIV-1 mAb have been determined (13, 29). This is also the case for the CH31 bNAb (11, 30). The antigen binding characteristic of the CH31 bNAb against a panel of HIV-1 Env antigens has been determined, including monovalent forms of the Env protein (CH505TF, 426c, and A244) or smaller fragments carrying only the engineered outer domain (eOD) (31, 32) such as eODGT6 and eODGT8 (Table 1). From these HIV-1 antigens, the CH31 bNAb has the highest affinity towards A244 and binds the eODGT8 with the highest on rate. In our study of antigen binding to the B cell surface, we exposed IgM-BCR or IgG1-BCR Ramos B cells to either a monomeric or a tetrameric form of the five indicated Env antigens. Afterward, the antigen binding was determined by flow cytometry. The data obtained are presented in Figure 3 according to the increased affinity of the CH31 bNAb in binding the 5 selected HIV-1 antigens.

**Table 1.** Env-derived antigens tested in the current study.

| Affinities and binding kinetics of HIV antigens to CH31 IgG1 antibody |  |  |  |  |
| --- | --- | --- | --- | --- |
| Ag | K<sub>D</sub> (nM) | k<sub>a</sub> (1/Ms) | k<sub>d</sub> (1/s) | Size KDa |
| eODGT6 (GT6) | **5.2μM** | 4.0E+4 | 2.1E-1 | 23,7 |
| CH505TF gp120 (CH505TF) | **314.2** | 1.2E+4 | 3.7E-3 | 53,1 |
| eODGT8 (GT8) | **307.8** | 2.3E+5 | 7.0E-2 | 22,9 |
| 426c core (426c) | **108.8** | 2.7E+3 | 3.0E-4 | 40,2 |
| A244 gp120 (A244) | **30.5** | 2.4E+3 | 7.3E-5 | 54,8 |

**Figure 3.**
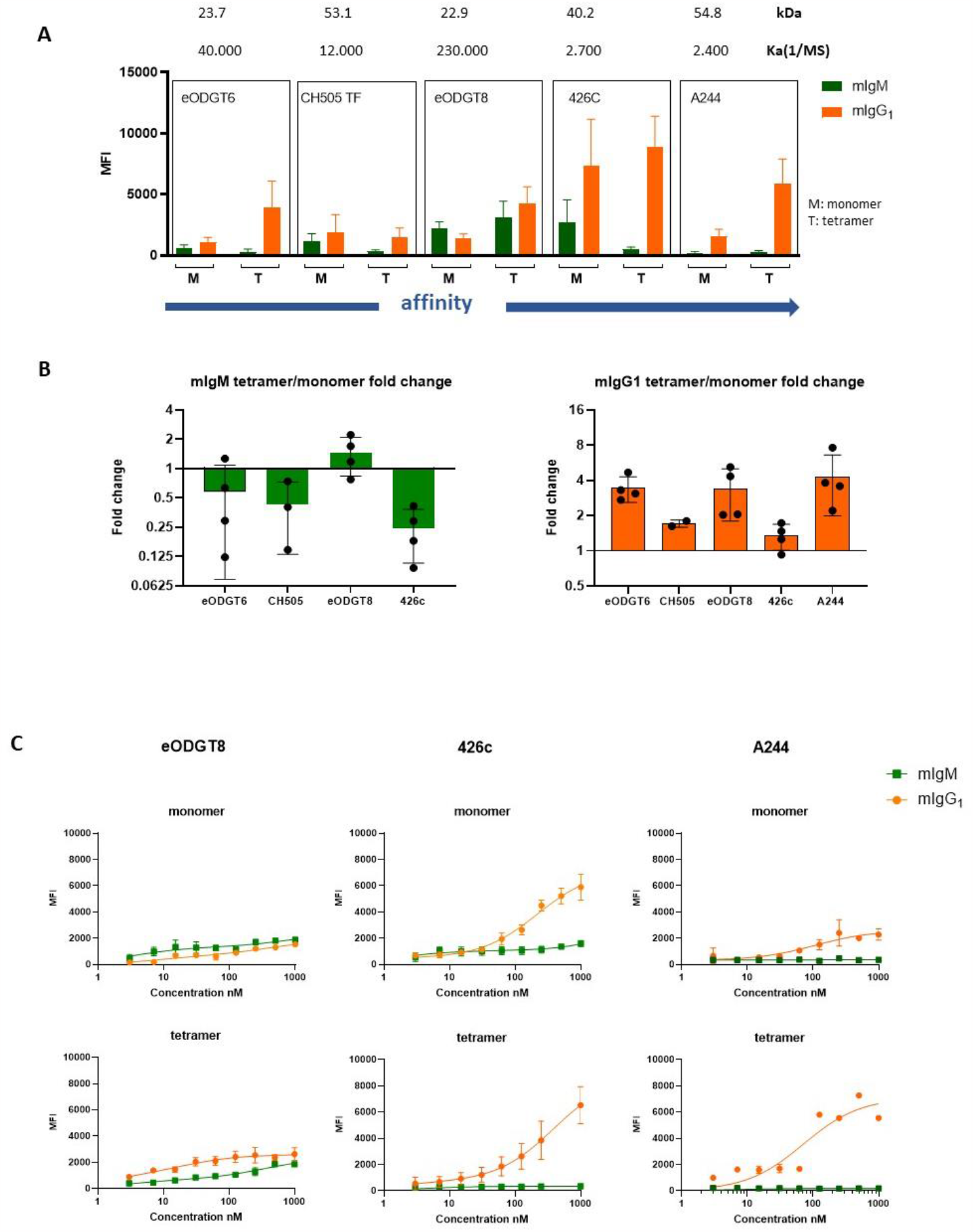
Monomeric and tetrameric antigen binding to CH31 IgM-BCR and IgG1-BCR cells. **(A)** monomeric and tetrameric forms of the Env antigens bound to IgM-BCR and IgG_1_-BCR. Cells were exposed to 1μM of each antigen for 30 min at 4°C, and thereafter the amount of antigen bound was detected with fluorescent probes and measured by flow cytometry; shown are the MFI values and SD (4 experiments were done per cell line). **(B)** Fold-change analysis of the binding of the tetramer compared to the monomer (MFI tetramer divided by MFI monomer) showing different binding of monomers and tetramers by each BCR (Y axis, Log2). **(C)** Antigen binding curves for eODGT8, 426c, and A244 monomer and tetramer, cells were exposed to a concentration range of the antigens (3nM to 1uM) for 30 min at 4°C, then the amount of antigen bound to the cells was detected with fluorescent probes and analyzed by flow cytometry, (X-axis, concentration Log10).

Using a lentiviral transfection system, a CH31 IgM-BCR Ramos B cell line has been previously generated and studied (11). It was found that this cell line best senses the HIV-1 antigen eODGT8 that is bound by the CH31 bNAb with the highest on-rate. Our retrovirally transfected CH31 IgM-BCR Ramos B cell also shows a preference for the eODGT8 antigen (Figure 3A). In addition, we find that the CH31 IgM-BCR binds more strongly to monomeric than tetrameric forms of the majority of antigens under study except for eODGT8. The parallel study of CH31 IgG1-BCR Ramos B cells shows a drastically different antigen binding pattern. The CH31 IgG1-BCR binds preferentially to the tetrameric rather than to the monomeric form of the 5 HIV-1 antigens. It also binds well to the tetrameric form of the A244 HIV-1 antigen that is not bound by the CH31 IgM-BCR although it is the antigen that is bound by the soluble CH31 IgG mAb with the highest affinity (Figure 3A, 5th panel). The different HIV-1 antigen binding characteristic of the CH31 IgM-BCR or IgG1-BCR is best seen by the fold-change analysis of the binding of tetrameric versus monomeric forms of the HIV-1 antigens (Figure 3B). Thus, while the CH31 IgM-BCR bound best to a monomeric antigen that gave the fastest on-rate, the IgG1-BCR bound best to the tetrameric form of high affinity HIV spike antigens.

To test the dependence of the binding to the CH31 IgM-BCR or CH31 IgG1-BCR Ramos B cells on the dose of antigen we conducted a dose-response assay with the monomeric or tetrameric forms of the HIV-1 antigens eODGT8, 426c, and A244 (Figure 3C). In spite of a 5-fold lower BCR expression level, the CH31 IgM-BCR Ramos B cells bind better to the monomeric form of the outer domain antigen eODGT8 than the CH31 IgG1-BCR Ramos B cells which in turn bind better to eODGT8 tetramers at all doses tested. The CH31 IgG1-BCR Ramos cells display an increasing antigen binding when exposed to increasing doses of the monomeric or tetrameric forms of the antigens 426c and A244 whereas the CH31 IgM-BCR Ramos hardly bind to these antigens even at higher doses. These data suggest that the CH31 IgM-BCR has difficulties to approach the CD4bs epitope in the bulkier gp120 antigen. It is notable that the CH31 IgG1-BCR binds better to 426c than A244 Env. The former antigen is lacking several glycan groups to overcome the steric hindrance encountered by the VRC01 bNAb family (33). In summary, these studies show a class-specific binding of HIV-1 antigens by the IgM-BCR and IgG1-BCR.

### The antigen binding characteristics of the CH31 IgG1-BCR can be transferred with the CH1γ domain

Different classes of the BCR can reside inside distinct protein/lipid nanodomains on the B cell plasma membrane and this could influence their antigen binding and signaling behavior (24). We thus first tried to disturb this organization by exposing the Ramos B cells with a panel of mAb against BCR coreceptors without changing the class-specific antigen binding (data not shown). We then focused our attention on the mIg structure and expressed mIgM/mIgG chimeric molecules on the Ramos B cell surface. The most informative of these chimeric constructs was one with an exchange of the CH1 domain of mIgM by that of the mIgG1 molecule (Figure 4A). The chimeric CH31 IgM(CH1γ)-BCR is highly expressed on the Ramos B cell surface and also carries significantly higher amounts of the B cell coreceptors and regulator proteins CD19 and CD45R (Figure 4B). Thus, the higher expression of the IgG1-BCR and coreceptors on the Ramos B cell surface could be transferred to the IgM-BCR with the CH1γ domain. A barcoded staining experiment with our panel of 29 mAb directed against different B cell surface markers also shows that the phenotype of the chimeric IgM(CH1γ)-BCR expressing Ramos B cells resembles more that of the IgG1-BCR-than the IgM-BCR Ramos B cells. (Supplementary Figure 1).

**Figure 4.**
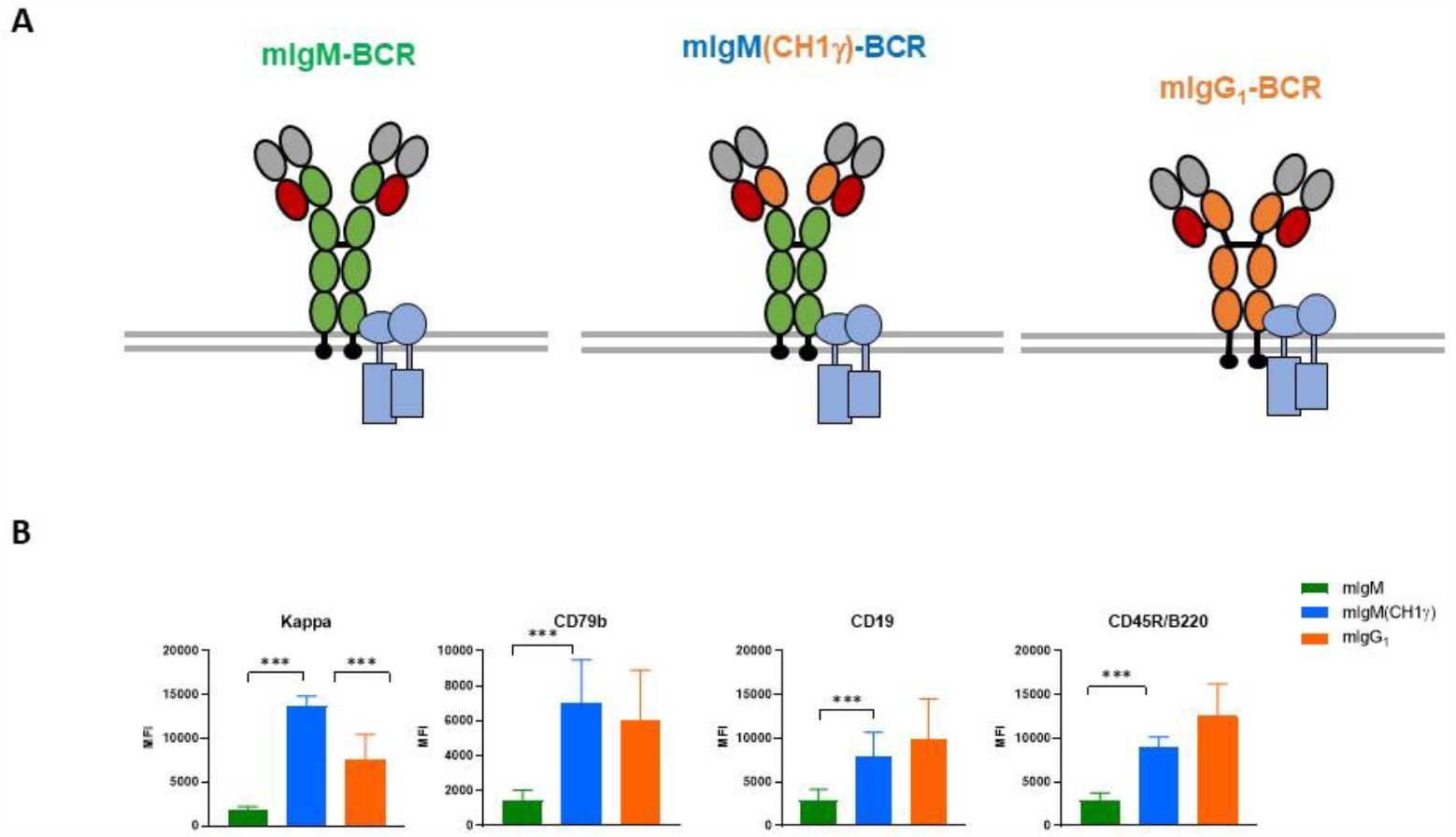
Role of CH1 domain on BCR expression. **(A)** Schematic drawing design of IgM chimeric BCR carrying the first constant domain of IgG1. **(B)** BCR components expression on IgM-BCR (n=16), IgG1-BCR (n=12), and the chimeric IgM(CH1γ)-BCR (n=5) cell lines, were evaluated by flow cytometry and expressed as MFIs and SD. Mann-Whitney test, *** p<0.001.

We next compared the antigen binding behavior of the chimeric CH31 IgM(CH1γ)-BCR (Figure 5A). Similar to what was found for the CH31 IgG1-BCR the chimeric CH31 IgM(CH1γ)-BCR binds more efficiently to the tetrameric than to the monomeric form of the HIV-1 antigens. In the case of the tetrameric CH505TF antigen, the chimeric BCR has a significantly higher binding efficiency than the CH31 IgG1-BCR. This binding preference of the CH31 IgM(CH1γ)-BCR for tetrameric HIV-1 antigens is clearly seen by the fold-change analysis of binding (Figure 5B). Thus, the replacement of the mIgM CH1 domains by that of the mIgG1 molecule drastically alters the expression level and the binding characteristic of the CH31 IgM-BCR.

**Figure 5.**
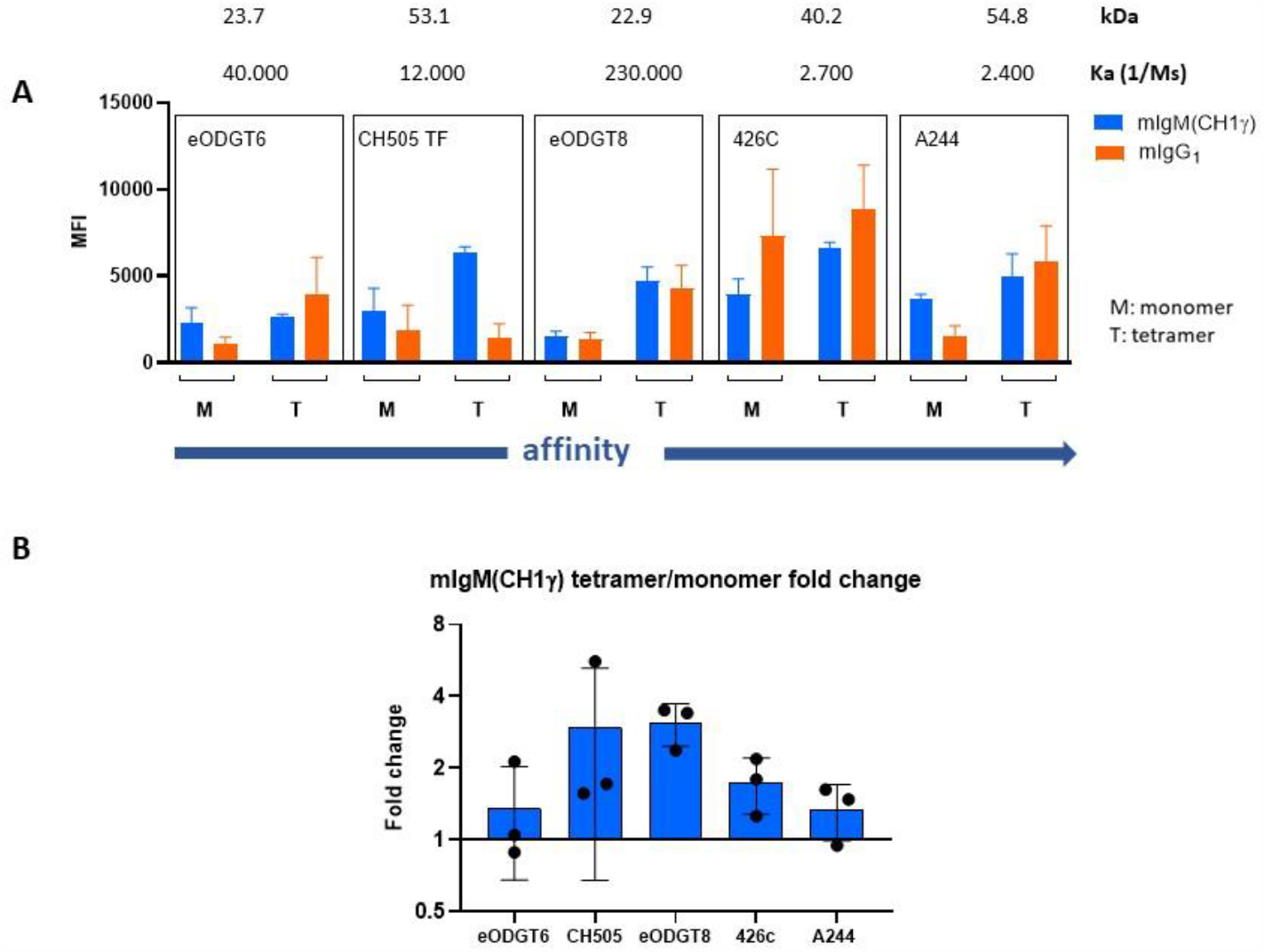
Role of CH1 domain on antigen binding. **(A)** Chimeric IgM(CH1γ)-BCR binding to monomer and tetramer Env antigens, cells were exposed to 1μM of each antigen for 30 min at 4°C, the amount of antigen bound was detected with fluorescent probes, shown are the MFI values and SD, the IgG1 antigen binding is shown for comparison purposes. **(B)** Fold-change analysis of antigen binding (MFI tetramer divided by MFI monomer, Y axis Log2).

### Antigen-induced calcium mobilization of BCR-positive Ramos cells

We next evaluated the antigen-dependent activation of the HIV-1 Env-specific Ramos B cells by a calcium mobilization assay (Figure 6). As previously reported (11), none of the monomeric forms of the 5 HIV-1 antigens under study induce a calcium flux response in the CH31 IgM-BCR Ramos cells and the same is true for the CH31 IgM(CH1γ)-BCR or CH31 IgG1-BCR Ramos B cells (data not shown). A calcium flux can be induced in all three Ramos lines after their exposure to 10 μg/ml of anti-human κLC antibodies. The reduced response of the CH31 IgM-BCR Ramos cells in this assay may be due to the lower BCR expression of these cells. The CH31 IgM-BCR Ramos cells do not show any response upon exposure to the CH505TF SOSIP trimeric HIV-1 antigen that induces a strong calcium flux in the antigen-stimulated chimeric CH31 IgM(CH1γ)-BCR and IgG1-BCR Ramos cells. However, all three studied Ramos cell lines display the calcium flux response when exposed to the high valency ODGT8 60mer. In summary, the Ramos cells expressing the chimeric CH31 IgM(CH1γ)-BCR and the CH31 IgG1-BCR not only bind more efficiently high valency HIV-1 antigens than CH31 IgM-BCR Ramos cells but react upon exposure to HIV-1 antigens with a stronger calcium flux.

**Figure 6.**
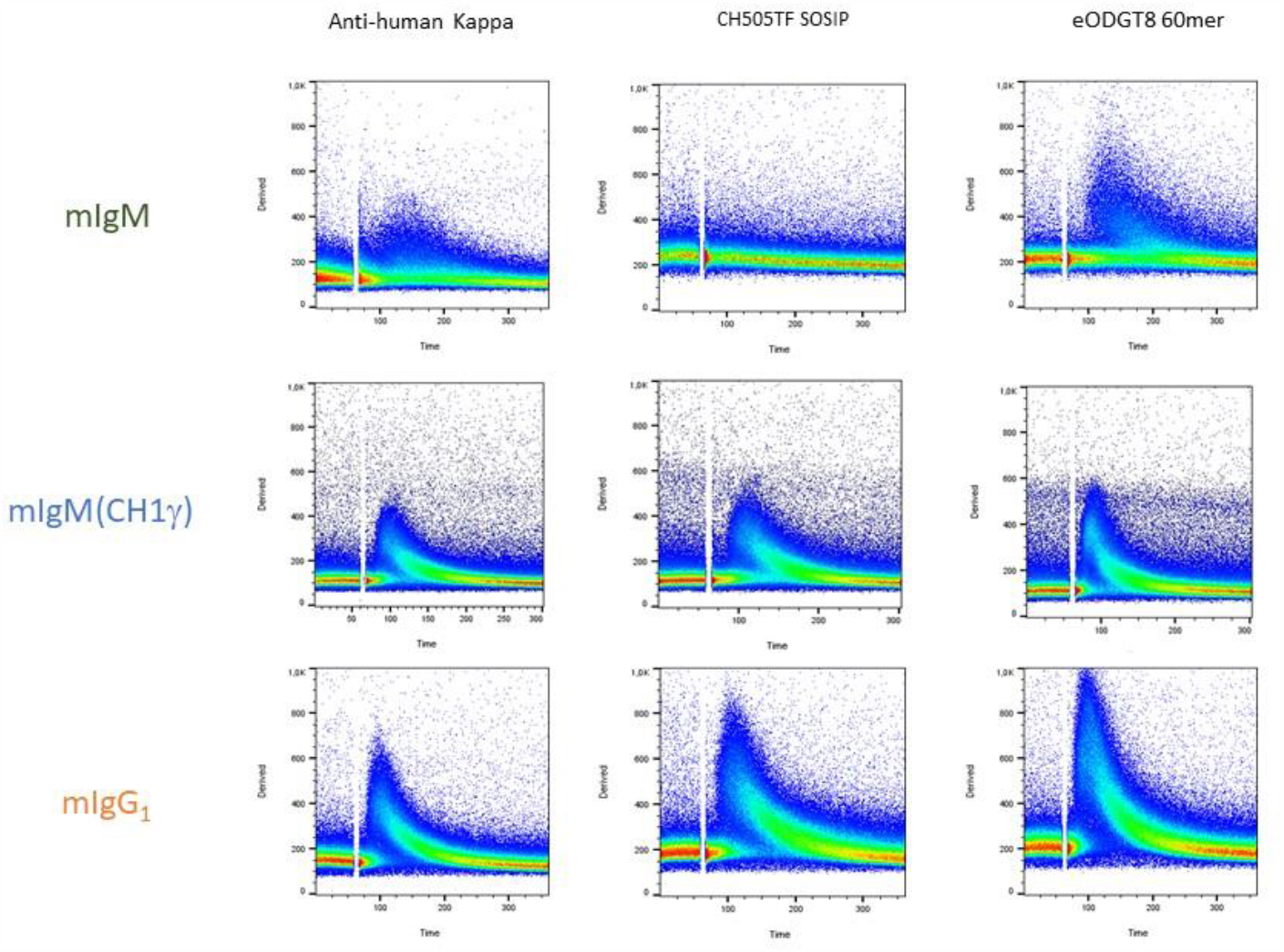
Calcium flux response to multimeric antigens. Calcium mobilization response of 1X10^6^ IgM-BCR, IgM(CH1γ)-BCR, and IgG1-BCR cells to stimulation with anti-human κ antibody compared to the stimulation with eODGT8 60 nanoparticle (200nM) and CH505 TFv4.1 SOSIP trimer (700nM), Results representative of duplicate experiments

**Figure 7.**
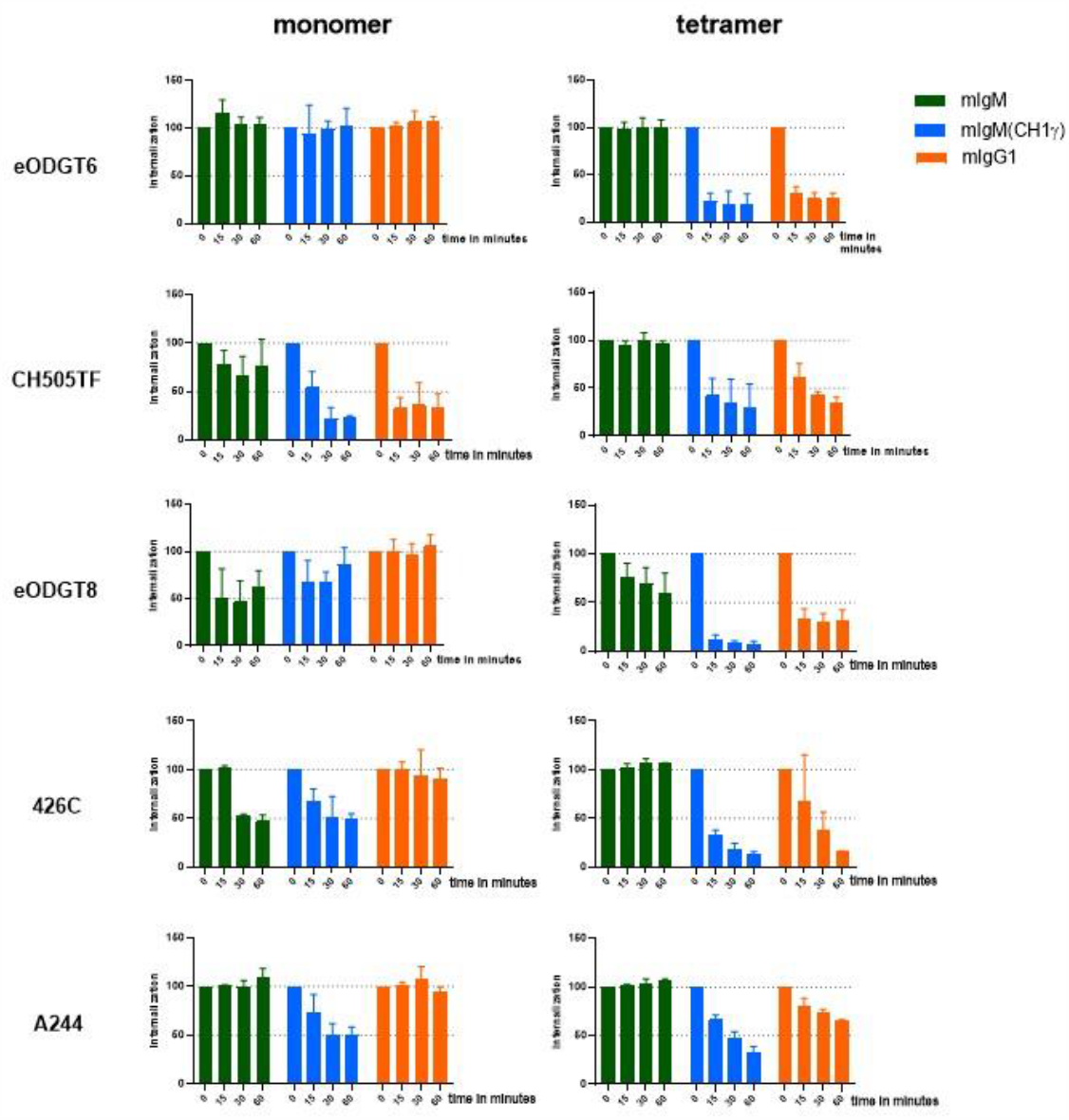
Antigen internalization in IgM-BCR, IgM (CH1γ)-BCR, and IgG1-BCR cell lines. Results show the monomer and tetramer antigen-induced internalization. The IgM-BCR, IgM(CH1γ)-BCR, and IgG1-BCR cell lines were separately exposed to 1uM concentration of each of the antigens after the initial binding incubation for 30 min at 4°C cells were moved to 37 degrees for different time points (see materials and methods). The changes in the amount of antigen bound by each BCR during the course of 60 min were measured by flow cytometry, and the MFIs from each time point were normalized to the unstimulated cells in each cell line. Results are shown as mean and SD of at least 3 experiments for each cell line.

### BCR internalization in response to antigen stimulation on IgM, IgG, and chimeric cells

The two major function of the BCR during antigen sensing is the activation of intracellular signaling pathways and the internalization of bound antigens leading to their processing and loading of antigenic peptides onto major histocompatibility complex (MHC) II proteins that promote T: B cell collaboration and full B cell activation (34). We thus determined the time course of antigen internalization of the CH31 IgM-BCR, IgM(CH1γ)-BCR, or IgG1-BCR Ramos B cells exposed for different times (15, 30, and 60 minutes) to 1μM of the monomeric or tetrameric forms of the five HIV-1 antigens under study (Figure 8). The internalization of monomeric HIV-1 antigens correlated well with their binding to the CH31 IgM-BCR Ramos cells (Figure 3). Only three (CH505TF, eODGT8, and 426) of the five antigens under study were thus internalized to some degree by these cells. In contrast, the internalization of monomeric HIV-1 antigens by the CH31 IgG1-BCR Ramos cells did not correlate well with their antigen-binding efficiency. For example, the monomeric 426 is well bound but is hardly internalized by the CH31 IgG1-BCR Ramos cells. From the 5 monomeric HIV-1 antigens, only the CH505TF was efficiently internalized although it showed a moderate binding to the CH31 IgG1-BCR Ramos cells. Interestingly the Ramos cells expressing the chimeric CH31 IgM(CH1γ)-BCR show some features of the CH31 IgM-BCR Ramos cells. Four of the five monomeric HIV-1 antigens under study were partly internalized by the CH31 IgM(CH1γ)-BCR Ramos cell in good correlation to their antigen binding efficiency.

All tetrameric forms of the five HIV-1 antigens under study are well internalized by both the CH31 IgM(CH1γ)-BCR and CH31 IgG1-BCR Ramos cells. Thus, in its sensing of tetrameric HIV-1 antigens, the chimeric CH31 IgM(CH1γ)-BCR behaves nearly identically to the CH31 IgG1-BCR. In contrast, the CH31 IgM-BCR Ramos internalized only one of the five tetrameric HIV-1 antigens namely eODGT8 which as a monomer is bound by the CH31 bNAb with the highest on-rate. In summary, we found that the binding of the diverse HIV-1 antigens is not only determined by the paratope formed by the VH/VL combination but also by the class of the CH1 domain.

## DISCUSSION

We here investigated the impact of the mIg class on the sensing of HIV-1 antigens. For this, we employed Ramos B cells expressing different classes of the HIV-1-specific CH31 BCR and a panel of HIV-1 antigens that vary in their size and binding kinetics. Antigen sensing studies cannot easily be conducted with human B lymphocytes as most of them carry BCRs of unknown antigen specificity. The germinal center-derived Burkitt lymphoma B cell line Ramos is a useful system for learning more about human B cell biology (23, 35, 36). These B cells have been frequently used for signal transduction studies (37) and their transcriptome, proteome, and surfacesome have been determined (38–40). Furthermore, with the CRISPR/Cas-9 technology, it is now straightforward to introduce mutations or gene KOs in this cell line. In particular, our MDL-AID KO Ramos B cells have been used by several groups for the generation and study of antigen-specific human B cell lines ^22,23^. Here we have employed this system to generate HIV-1-specific B cells expressing either a CH31 IgM-BCR or a CH31 IgG1-BCR and found a 5-fold higher expression of the latter BCR class on the Ramos B cell surface. Interestingly, the high expression phenotype can be transferred to the IgM-BCR with the CH1γ domain. Unlike any other CH domain, the CH1 domain cannot fold autonomously in the endoplasmic reticulum (ER) but requires interaction with the CL domain for its proper folding (41). The partially unfolded state of the CH1 domain is detected by the ER quality control system. Specifically, the binding immunoglobulin protein (BiP), also known as heat shock protein family A member 5 (HSPA5) can directly bind to the partially folded CH1 domain and prevent the export of free immunoglobulin heavy chains. The CH1/BiP complex is not only responsible for the retention of the free HC in the ER but also promotes CH1/CL conjugation which is required for proper CH1 folding (42, 43). In comparison to the CH31 IgM-BCR, the higher expression of the chimeric CH31 IgM(CH1γ)-BCR may be due to a more efficient folding and/or binding of the CH1γ to the CL domain. In favor of the latter possibility is the high sequence conservation of the 3 beta strands connecting the CH1γ to the CL domain. The IgM-BCR Ramos cells express more BiP than the IgG1-BCR Ramos cells (unpublished observation), suggesting that the former B cells have a higher level of unfolded CH1 domains and free HCs. The higher expression of B cell surface markers on the IgG1-BCR or IgM(CH1γ)-BCR than on IgM-BCR Ramos cells maybe do to the co-localization of these markers with the BCR. This is the case for CD45 which also interacts with the tetraspanin CD53 on the lymphocyte membrane (44). Furthermore, the chemokine receptor CXCR4 is functionally associated with the BCR on naïve murine B cells (45, 46).

Despite their identical VH/VL domains, the different classes of the CH31 BCR have distinct antigen sensing capabilities in that the CH31 IgM-BCR and CH31 IgG1-BCR preferentially bind monovalent and polyvalent (tetrameric) antigens, respectively. A major finding of our study is that this class-specific antigen-sensing behavior can be transferred with the CH1γ domain from the IgG-BCR to the IgM-BCR. Our studies support the notion that the CH1 domain not only plays an important role in the assembly of mIg molecules but also in the sensing of cognate antigens. The influence of the CH1 domain on antigen binding has previously been noticed in studies of antigen-specific mAb. For example, Pritsch et al. (47) described in 1996 two different classes of anti-tubulin mAb with identical VH and VL sequences and showed that the IgA1 class binds tubulin with a ten times higher affinity than the IgG1 class mAb. These different binding features were also found in Fab fragments of these antibodies, suggesting that it is the CH1 domain that determines the class-specific binding kinetics. In the meantime, the class-specific antigen binding of mAb with identical VH and VL sequence has been verified by several studies of different antigenic systems and have been summarized in a recent review (48). Interestingly, this summary shows that the class-specific antigen sensing of a mAb is dependent on the VH family used and on the LC isotype. Specifically, there are permissive and non-permissive VH family members, and only mAb carrying a κLC but not those with a λLC are showing this feature. In line with this classification is the observation that the CH31 BCR carries a permissive VH1-2 domain and a κLC. Our study is the first to demonstrate that class-specific antigen sensing is not only a feature of soluble mAbs but also of membrane-bound antigen receptors and influences not only antigen binding but also receptor signaling and internalization.

An important aspect of our study is that we could directly compare the class-specific sensing capabilities of CH31 IgM-BCR and CH31 IgG1-BCR B cells exposed to HIV-1 antigens of different sizes and binding kinetics. In connection to the IgM class, the CH31 paratope seems to be a more rigid structure that is unable to adapt to the larger HIV-1 antigens 426c and A244 although they are bound with the highest affinity by IgG-class CH31 bNAb. As reported previously (11), the CH31 IgM-BCR binds most efficiently to the smaller and truncated eODGT8 antigen which has the largest on-rate of binding and thus seems to interact with a preformed part of this rigid paratope. Upon transfer of the CH1γ domain, the IgM-BCR gains the binding characteristic of the IgG1-BCR and prefers larger and polymeric HIV-1 antigens. Thus, the CH1 has a major impact on the rigidity or flexibility of the CH31 paratope.

Currently, there are two non-exclusive possibilities for how the CH1 domain could have an impact on antigen sensing. One is that a permissive VH domain is connected to the CH1 domain via specific amino acids in the opposing Ig-domain loops at the VH/CH1 interface. Although the existing crystal structures of mAbs do not show such an interdomain amino acids conjugation, the VH/CH1 interface interactions may be highly dynamic and thus missed by crystallographic techniques (11). In line with this possibility are studies showing antigen-induced conformational changes in the CH1 domain loops that may influence the orientation of the two Fab arms of a mAb (49). If, indeed, antigen-binding can alter the CH1 domain conformation then the different classes of the CH1 domain may have an impact on the orientation of the CDR loops of the VH domain and thus on antigen binding.

Alternatively, the class of the CH1 domain could influence the binding character of a mAb or BCR via its connection to the CL domain. For example, it has been found that Fab fragments of antibodies can display different angles between the HC and the LC part and that these different angles could influence the antigen sensing behavior (50). The finding that only the κ but not the λ LC is associated with class-specific antigen sensing (48) suggests that the Cλ domains domain has an impact on the function of the CH1 in antigen sensing. The Cλ domain may be more rigid than the Cκ domain and thus less flexible to adopt different Fab structures that have an impact on antigen sensing.

Understanding the molecular features that account for the differences in antigen sensing of an IgM-BCR or IgG-BCR can have an impact on the design of optimal vaccines and immunization protocols. Currently, the same vaccine formula is given for the primary and the following (booster) vaccinations. Our finding that the CH31 IgM-BCR preferentially binds monovalent, whereas the CH31 IgG-BCR binds polymeric (tetrameric) antigens suggests that vaccination protocols should be modified. A primary vaccine should mostly consist of monomeric proteins, whereas for the secondary booster response multimeric formulas of antigens should be given. One caveat of our study, however, is that our IgM-BCR Ramos cells are different from naïve B cells as they express only one BCR class, whereas naïve B cells co-express the IgD-BCR and IgM-BCR. Recent data show that, unlike the IgM-BCR, the IgD-BCR cannot sense monovalent antigens (51) and it is therefore possible that naïve B cells gain their ability to sense polymeric antigens via the IgD-BCR. Nevertheless, it will be interesting to see whether the combination of monovalent primary vaccines with polymeric secondary vaccines has a different impact compared to the current vaccination protocols.

## MATERIALS AND METHODS

### Cells

Human Ramos MDL AID KO (HC, LC, activation-induced cytidine deaminase KO) (23), and Phoenix packaging cell line were cultured in RPMI medium (Gibco) supplemented with 10% Fetal calf serum (FCS, PAN Biotech), 10 units/mL penicillin/ streptomycin (Gibco) and 50 mM β-mercaptoethanol (Sigma). The CH31 IgM-BCR, IgG1-BCR, and IgM(Cγ1)-BCR Ramos B cells were cultured in RPMI 5% FCS. All cell lines were incubated at 37°C with 5% CO2 and split every 2 days.

### Cloning

For the retroviral expression vectors, sequences encoding the mIgM, mIgG1, and chimeric mIgM(CH1γ) with the CH31 bnAb specificity, were cloned on the pMIG vector backbone (pMIG Addgene plasmid # 9044). The cDNA coding for the VH and VL domains of CH31 was produced by Integrated DNA Technologies, IDT (Leuven Belgium) as gBlocks, and cloned into the pMIG vector by In-Fusion cloning (ClonTech, Takara). For the design of the cloning primers and plasmids, Geneious 9.0.5 software was used. The components of the master mix for the PCR and the PCR programs were set up according to the CloneAmp HIFI PCR protocol (Takara). All generated plasmids were sequenced before use (Eurofins Genomics).

### Retroviral transfection in Phoenix Cells and transduction of Ramos Cells

Phoenix cells were split every 2 days by diluting them 1/10. Twenty-four hours before the transfection, 5x10^5^ Phoenix cells were plated into a 6-well plate in 2 ml of RPMI 10%. The transfection was performed on cells at 70 % confluency, according to the manufacturer’s instructions, using Polyjet transfection reagent, retrovirus-producing plasmids pKAT, and pMIG plasmid carrying the mIg coding sequences. After 2 days of culture, the virus-containing supernatant was collected and filtered through a 0.45 μm filter, afterward, Polybrene was added to the viral supernatant at a concentration of 1 μl/ml. Ramos MDL AID KO cells were split the day before the transduction, 5x10^5^ Ramos cells were resuspended in 1 ml of the transduction mixture, plated in 12-well plates, and incubated overnight at 37°C. Cells were then washed and resuspended in RPMI 10% FCS. Transfection efficiency was evaluated after 48 hours.

### Antibodies

A list of the antibodies used for the phenotyping is included in a supplementary table.

### Barcode phenotype analysis

To phenotype all the CH31 Ramos B cell lines simultaneously and with accuracy, barcoding and surface staining techniques were used. The barcoding involved preparing different serial concentrations of the cell proliferation tracers in PBS, CytoTell blue or CytoTell green (both form AAT BioQuest). The Ramos B cells were then resuspended in a mixture 1:1 of both tracers and incubated in the dark for 20 minutes at room temperature. After washing the cells with PBS, they were combined in one test tube for further analysis. For surface staining, the barcoded cells were stained with antibodies in FACS Buffer (PBS, 3% FCS, 0.1% sodium azide), incubated, and measured using an Attune Nxt Acoustic Focusing Cytometer. This process ensured reliable and reproducible results. Data was analyzed with FlowJo software (BD Biosciences). Shown are the median fluorescence intensity (MFI) and standard deviation (SD) of IgM-BCR n=16, IgG1-BCR n=12, and IgM(CH1γ)-BCR n=5 repetitions. Analysis was done using GraphPad Prism 9.5.1

### Antigens

The following antigens were produced at the Duke Human Vaccine Institute (11): eODGT6 (KX527852), eODGT8 (KX527855), 426c core WT (KX518323), A244 D11gp120, CH0505-Con D7 gp120, CH505TFv4.1 SOSIP. The monomer form of the antigens was biotinylated and the subsequent tetramerization of these proteins was done with FITC-labeled avidin (Invitrogen, A821), in brief, monomeric antigens were incubated with a 5-fold molar excess of avidin and incubated for 1,5 h at room temperature, 900 rpm in a Thermomixer. The FITC molecule was used to detect tetramer binding and internalization in Ramos cells, using an anti-FITC antibody. eODGT8 d41m3 60mer (KX527857) was produced at The Scripps Research Institute.

### Antigen tetramerization

Tetramerization of monomeric and trimeric proteins was accomplished with Avidin-FITC (Invitrogen). A 4:1 molar ratio of protein to avidin was used to maximize avidin site occupancy; avidin was added stepwise to the protein. The appropriate volume of avidin was added every 15 min followed by agitation at 900rpm at room temperature. The final molarity of the protein was calculated based on the number of moles used in the reaction as well as the total volume of protein and avidin combined.

### Antigen binding to CH31 Ramos cells

The binding of Env antigens to the different classes of the CH31 BCR was evaluated using flow cytometry. For this purpose, 2.5x10^5^ cells were incubated with 1μM of different Env HIV-1 antigens in PBS 2% FCS, and incubated for 30 min at 4°C. Afterward, cells were washed and stained with a Neutravidin DyLight 633 (Invitrogen, 22844) to detect monomeric biotinylated antigens and anti-FITC antibody (Invitrogen) to detect Avidin-FITC conjugated tetrameric antigens. After incubation, cells were washed and resuspended in FACS Buffer (PBS, FCS 3%, and 0.05% azide) and analyzed in Attune Nxt Acoustic Focusing Cytometer. The results show the median fluorescence intensity (MFI) and standard deviation (SD). Analysis was done using GraphPad Prism 9.5.1

### Calcium measurements

Calcium measurements were performed as previously described (52). In brief, 1 × 10^6^ cells were loaded with Indo-1 AM (Molecular Probes) 5μg/ml, and puronic F127 (Molecular Probes) 5μg/ml, for 45 minutes at 37°C, following the manufacturer’
ss instructions. After incubation, cells were washed and resuspended in 500 μl RPMI medium supplemented with 1% FCS. Before analysis, the cells were pre-warmed at 37°C for 5 min. Stimuli were added after 1 minute of baseline recording. The calcium flux was measured with a Fortessa II (BD). Data was analyzed with FlowJo software (BD Biosciences).

### Antigen internalization assay

In this assay, we monitor antigens internalization by staining for HIV antigens that remain bound to CH31 BCR after different times of culture of the CH31 Ramos B cells. Cells were counted and washed once with PBS, and 1x10^6^ cells in 500 μl PBS 2% FCS were cooled for 10 min on ice, afterward, 1μM of antigen or a similar volume of PBS (negative control) was added and binding was allowed incubating 30 min on ice. Cells were washed once with ice-cold PBS 2 % FCS and resuspended in 500 μl. To allow internalization, aliquots of 2x10^5^ cells in 100 μl PBS 2 % FCS were prepared and incubated for 0, 15, 30, and 60 min at 37°C in a thermomixer (Eppendorf). Immediately after the incubation, BCR trafficking was stopped by placing the samples on ice and by adding 1 ml ice-cold FACS buffer. Cells were pelleted by centrifugation and resuspended in 100 μl staining solution; in order to detect antigens remaining on the surface, staining was done for 15 min on ice and in the dark. Subsequently, cells were washed twice with ice-cold FACS buffer and acquired at the Attune Nxt Acoustic Focusing Cytometer. Analysis was done using GraphPad Prism 9.5.1

## ACKNOWLEDGMENTS

We thank Frauke Bartels-Burgahn for her help with the cell cultures and Kathrin Klaesener and Dr. Lise Leclercq for reading and correcting the manuscript. We thank Dr. Barton F. Haynes, DHVI, Director of CHAVD (Duke Consortia) for providing facility resources. We are grateful to Kevin Saunders, and Elizabeth Donahue (Duke HVI) for expressing and purifying Env gp120 and gp140 trimers and NPs. We thank William Schief, and Bettina Groschel at Scripps Research for eODGT6 and eODGT8 proteins and eODGT8 60-mer nanoparticles.

## Data and materials availability

All data are available in the main text or the supplementary materials.

