## Supplementary material for "Class-specific sensing of HIV-1 antigens by the B cell antigen receptor depends on the CH1 domain"

**SUPPLEMENTARY MATERIALS**

**Supplementary Figure 1**. Phenotype differences among CH31 IgM (CH1γ)-BCR chimeric BCR and the CH31-IgM BCR and CH31 IgG-BCR Ramos cells. Volcano plot depicting differentially expressed surface markers of the CH31 IgM (CH1γ)-BCR when compared to IgM-BCR (green plot) and IgG BCR cells (orange plot), shown are the mean values of IgM n=16, IgG n=12, and IgM (CH1g) n=5 repetitions. Analysis was done using GraphPad Prism 9.5.1.

**Supplementary Figure 2. A.** Schematic representation of the CH31 IgM (CH1γ-L) Chimeric BCR, in which the CH1μ domain was replaced by the CH1γ domain and the hinge region of IgG. **B.** Antigen internalization, both chimeric cells CH31 IgM (CH1γ)-BCR and CH31 IgM (CH1γ-L)-BCR were stimulated with 1μM of the monomeric forms of the antigens and the changes in the amount of antigen bound by each BCR during the course of 60 minutes were measured by flow cytometry. The MFIs from each time point were normalized to the unstimulated cells in each cell line. Results are shown as mean and SD of at least 3 experiments for each cell line.

**Supplementary table 1.** List of the antibodies used to phenotype Ramos B cells

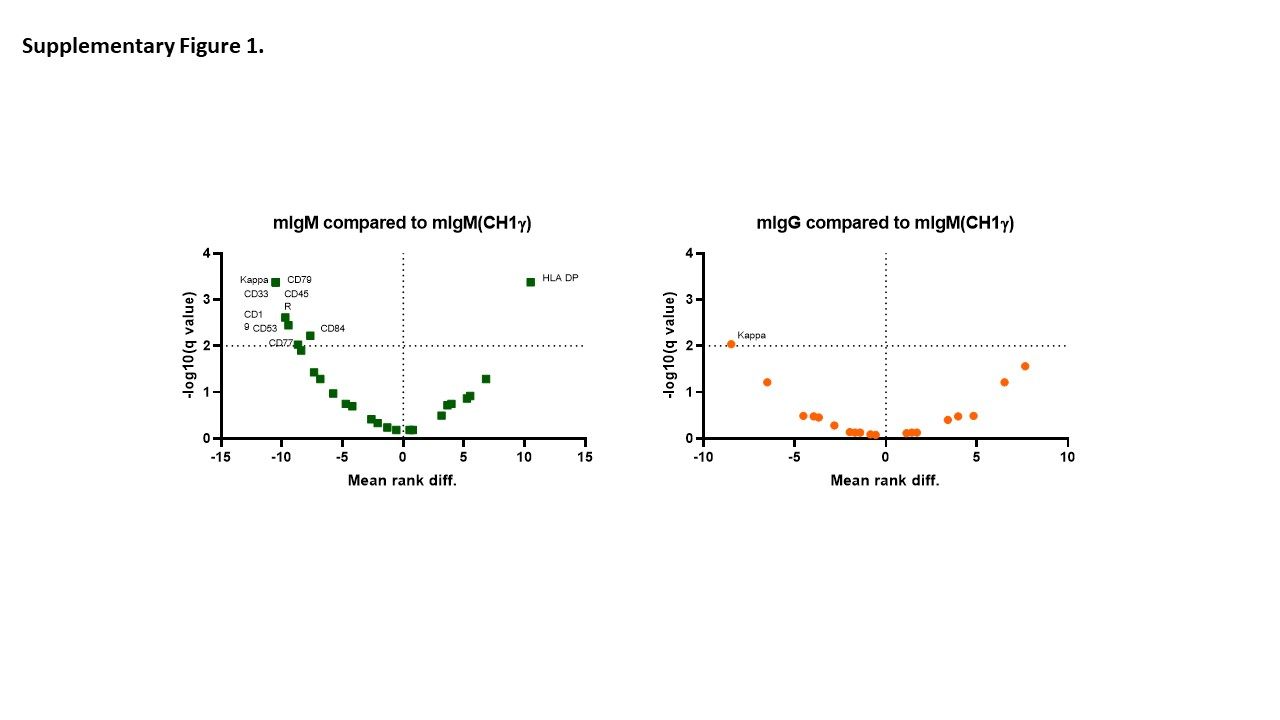

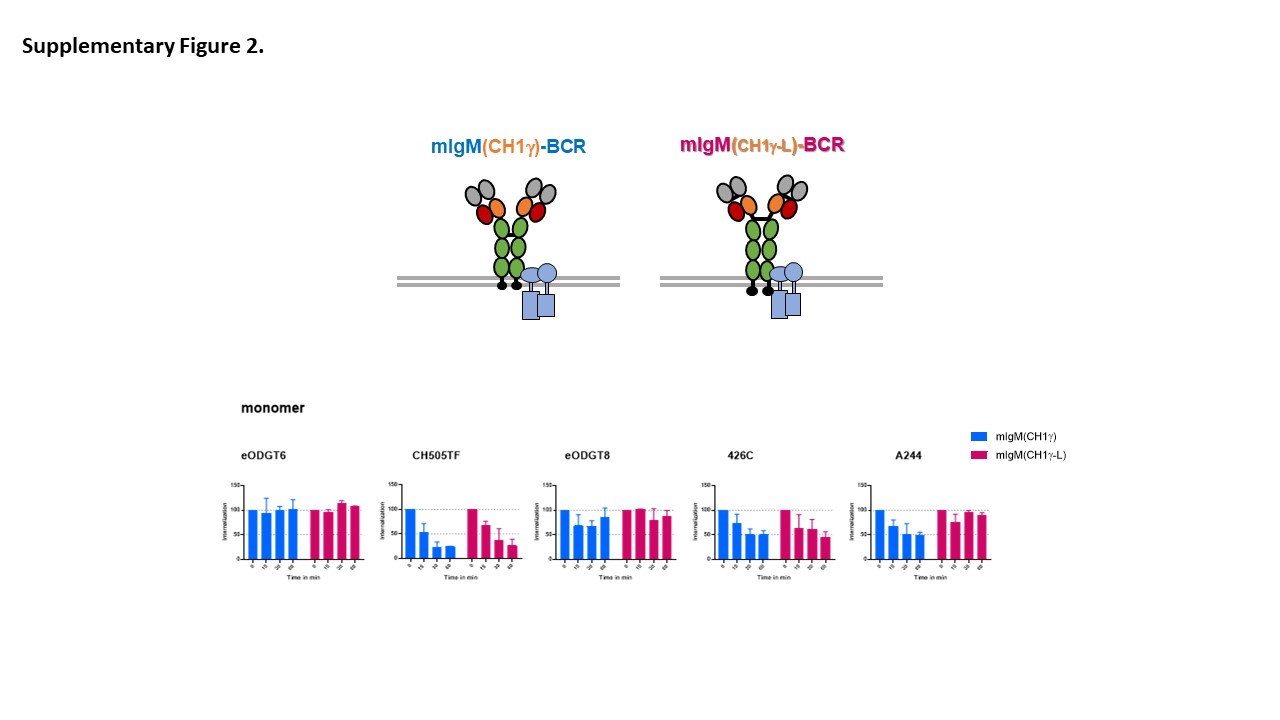

| **ANTIBODY** | **CATALOG** | **Clone** |
| --- | --- | --- |
| Anti human CD19 APC | BioLegend 302212 | HIB19 |
| Anti human CD25 PE | BD 555432 | MA251 |
| Anti human CD27 PECy7 | BioLegend 356412 | MT271 |
| Anti human CD37 AF647 | BD 561562 | MB371 |
| Anti human CD40 PE | BioLegend 334308 | 5C3 |
| Anti human CD45R PE (B220) | eBiosciences 12045285 | RA3-6B2 |
| Anti human CD53 PE | BioLegend 325406 | HI29 |
| Anti human CD63 PE | BioLegend 353004 | H5C6 |
| Anti human CD71 PE | BioLegend 334106 | Cy1G4 |
| Anti human CD74 PE | BioLegend 326808 | LN2 |
| Anti human CD81 APC | eBiosciences 170819-42 | 1D6-CD81 |
| CTB AF647 | Thermofisher C34778 |  |
| Anti human CXCR4 PE (CD184) | BD 555974 | 12G5 |
| Anti human HLA ABC PE | BioLegend 311406 | W6/32 |
| Anti human HLA DP PE | Leinco technologies H130 | B7/21 |
| Anti human HLA DR APC | BioLegend 307610 | L423 |
| Anti human IgM APC | BioLegend 314510 | MHM88 |
| Anti human CD20 PeCy7 | BioLegend 302312 | 2H7 |
| Anti human CD38PeCy7 | BioLegend 303516 | HIT2 |
| Anti human CD69 PECY7 | BioLegend 310912 | FN50 |
| Anti human CD83 PECY7 | BioLegend 305326 | HB15 |
| Anti human CD29 PECY7 | BioLegend 303026 | TS2/16 |
| Anti human CD47 PE | BioLegend 323108 | CC2C6 |
| Anti human CD77 AF647 | BD 563632 | 5B5 |
| Anti human CD22 PECY7 | BioLegend 302514 | HIB22 |
| Anti human IgB(79b) PE | BioLegend 341404 | CB3-1 |
| Anti human CD72 PE | BioLegend 316208 | 3F3 |
| Anti human CD33 PECY7 | BioLegend 366618 | P67.6 |
| Anti human IgG PECY7 | BioLegend 410722 | M1310G05 |
| Anti human CD352 PE | BioLegend 317208 | NT-7 |
| Anti human CD84 PE | BioLegend 326008 | CD84.1.21 |
| Anti human CD84 APC | BioLegend 326010 | CD84.1.22 |
| Anti human Light chain kappa PE | BioLegend 392704 | TB28-2 |
| Anti FITC APC | Invitrogen 17-7691-82 | NAWESLEE |
